# Visual processing and fold-change detection by the larva of the simple chordate *Ciona*

**DOI:** 10.1101/2020.06.13.150151

**Authors:** Cezar Borba, Matthew J. Kourakis, Shea Schwennicke, Lorena Brasnic, William C. Smith

## Abstract

Visual processing transforms the complex visual world into useful information. *Ciona*, a close relative of vertebrates, has one of the simplest nervous systems known, yet has a range of visuomotor behaviors. Among them are negative phototaxis and a looming-shadow response. These two behaviors are mediated by separate photoreceptor groups acting via distinct, but overlapping, neural circuits. We show here that processing circuits underlying both behaviors transform visual inputs to generate fold-change detection (FCD) outputs. In FCD, the response scales with the relative fold changes in input, but is invariant to the overall magnitude of the stimulus. Several different circuit architectures can generate FCD responses. Both the behavioral outputs and the putative circuitry for the two visuomotor behaviors point to them using different FCD circuits. Pharmacological treatment points to circuits in the posterior brain vesicle of *Ciona*, a region we speculate shares homology with the vertebrate midbrain, as important for FCD.

## Introduction

*Ciona* has served as a valuable model organism both because of its close evolutionary relationship to the vertebrates, and because of its genetic, embryonic and anatomical simplicity (Lemaire et al., 2008; Satoh, 1994, 2014). Phylogenetically, *Ciona* is a member of the chordate subphylum known as the tunicates. Collectively, the tunicates comprise the closest extant relatives of the vertebrates (Delsuc et al., 2006). The close relationship of the tunicates to the vertebrates is evident at all scales - from genomic to anatomical. Particularly striking is the *Ciona* tadpole larva, which highlights both these attributes: vertebrate-like anatomy and simplicity. In common with similarly-staged vertebrates, the *Ciona* larva features a notochord running the length of its muscular tail and a dorsal central nervous system (CNS) with a central ventricle. Despite this conserved chordate anatomy, *Ciona* larval organs are composed of very few cells: 40 notochord cells, 36 tail muscle cells, and ~180 neurons in the CNS (Nicol and Meinertzhagen, 1991; Satoh, 1994). The conserved chordate features of the *Ciona* larval CNS extends to anatomical subdivisions, with homologs of the vertebrate forebrain, midbrain-hindbrain junction, hindbrain and spinal cord [reviewed in (Hudson, 2016)]. The simplicity of the *Ciona* larval CNS has enabled the generation of a complete synaptic connectome by serial-section electron microscopy (Ryan et al., 2016).

Although small in cell numbers, the *Ciona* larval CNS supports a number of sensory systems that direct a range of complex behaviors. These behaviors include negative gravitaxis, mediated by the otolith organ, mechanosensation, mediated by peripheral touch receptors, and two distinct visuomotor behaviors mediated by ciliary photoreceptors that cluster into two functional groups in the ocellus organ (Bostwick et al., 2020; Horie et al., 2008; Kajiwara and Yoshida, 1985; Ryan et al., 2018; Salas et al., 2018; Svane and Young, 1989). Figure 1A shows a simplified *Ciona* larva with the visuomotor circuits highlighted. In the figure, neurons of the same class are grouped together, as are the synaptic connections between them. The first of the two photoreceptor clusters is the PR-I group, which is composed of 23 photoreceptors (Figure 1A). All but two of the PR-I photoreceptors are glutamatergic (Figure 1B). For the other two PR-I photoreceptors, one is GABAergic, and the other is dual glutamatergic/GABAergic (Kourakis et al., 2019). The PR-I photosensory system mediates negative phototaxis with the aid of an associated pigment cell (*pc* in Figure 1) that directionally shades the outer segments of the photoreceptors, allowing larvae to discern the direction of light as they perform casting swims (Salas et al., 2018). The second ocellus photoreceptor cluster, the PR-II group, is composed of seven photoreceptors and is located anterior to the PR-I group [Figure 1A; (Horie et al., 2008)]. The PR-II group does not have an associated pigment cell, and evokes swimming in response to dimming ambient light, most likely as a looming-shadow escape behavior (Salas et al., 2018). The PR-II group contains a mixture of GABAergic and dual glutamatergic/GABAergic photoreceptors (Figure 1B) (Kourakis et al., 2019).

**Figure 1.**
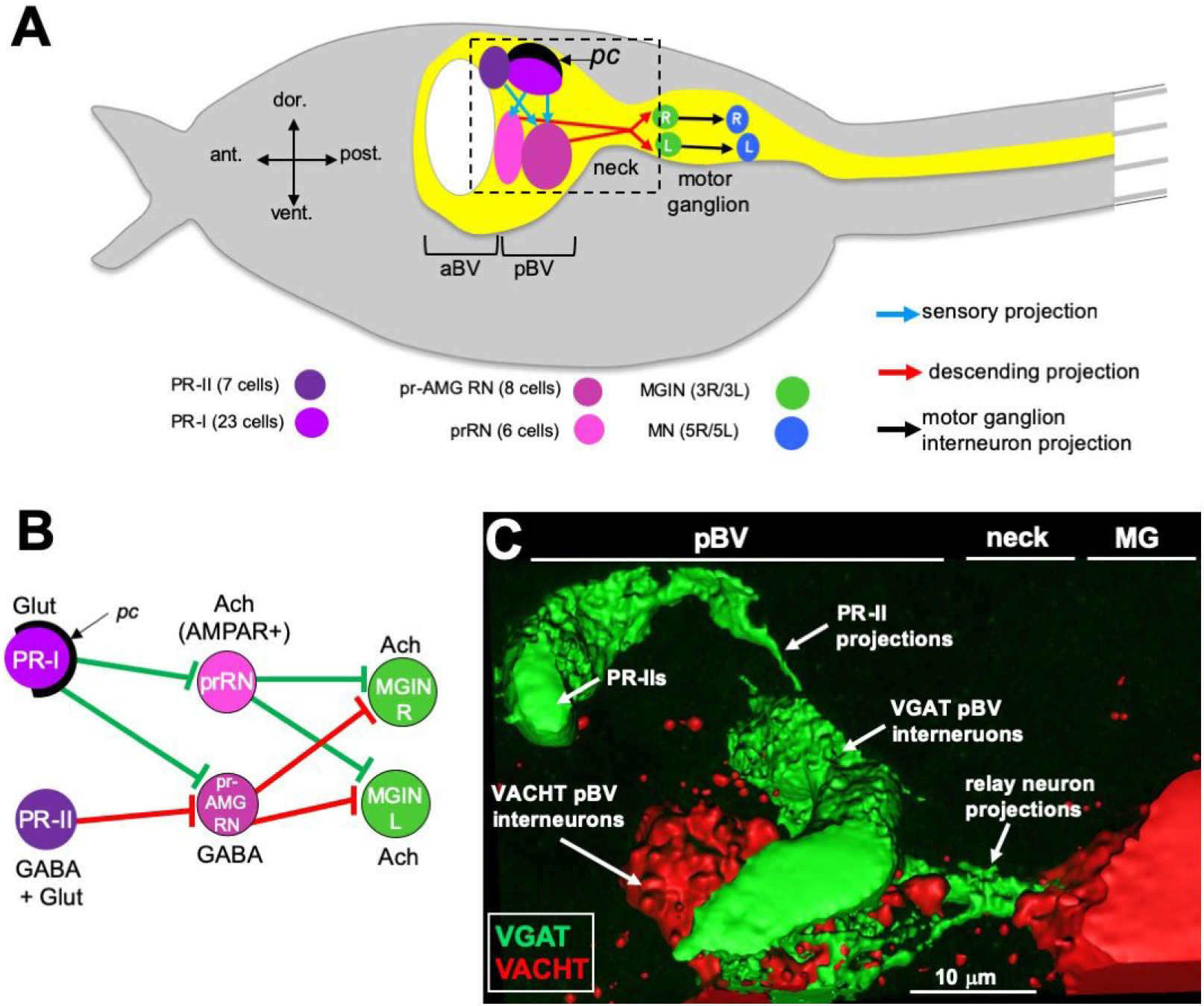
**A.** Cartoon of *Ciona* larva (only a portion of tail is shown). Highlighted in the CNS (yellow) are the minimal visuomotor pathways. Cell classes are color coded according to (Ryan and Meinertzhagen, 2019), and the number of cells in each class are indicated in parentheses. **B**. Minimal visuomotor circuits. Green lines indicate putative excitatory, and red lines putative inhibitory, synapses. **C**. GABAergic PR-II photoreceptors project to GABAergic relay neurons in the pBV. View corresponds approximately to the dashed box in panel A. Abbreviations: pc = pigment cell; aBV = anterior brain vesicle; pBV = posterior brain vesicle; PR = photoreceptor; pr-AMG RN = photoreceptor ascending motor ganglion relay neurons; prRN = photoreceptor relay neurons; MGIN = motor ganglion interneurons; MN = motor neurons; L = left; R = right; Glut = glutamate; Ach = acetylcholine; AMPAR = AMPA receptor; MG = motor ganglion; VGAT = Vesicular GABA Transporter; VACHT = Vesicular acetylcholine transporter.

Both the PR-I and PR-II photoreceptors project directly to relay interneurons in the posterior brain vesicle (pBV) (Figure 1A). These relay neurons in turn project primarily to the cholinergic *motor ganglion interneurons* (MGINs) of the motor ganglion (MG) (Figure 1A and B). pBV relay neurons with photoreceptor input fall into two main classes. The six *photoreceptor relay neurons* (prRNs) receive input from the PR-I group (Figure 1A and B). *In situ* hybridization studies indicate that the prRNs are predominantly, or perhaps exclusively, cholinergic (Kourakis et al., 2019). The other major class of pBV relay neurons with photoreceptor input are the eight *photoreceptor ascending motor ganglion relay neurons* (pr-AMG RNs; Figure 1A and B). The pr-AMG RNs are predominantly GABAergic and receive input from both the PR-I and PR-II groups (Figures 1A and B; Kourakis et al., 2019; Ryan et al., 2016). Thus the prRNs receive input only from the PR-Is, while the pr-AMG RNs receive input from both photoreceptor groups. Significantly, while both the pr-AMG RNs and the prRNs receive glutamatergic input from the PR-Is, only the cholinergic prRNs express the glutamate AMPA receptor [AMPAR; Figure 1B and (Kourakis et al., 2019)]. Moreover, treatment with the AMPAR antagonist perampanel blocks negative phototaxis while not disrupting the light-dimming response (Kourakis et al., 2019). Thus the minimal circuit for negative phototaxis appears to involve the glutamatergic photoreceptors stimulating the cholinergic prRNs, which then project to the MG to stimulate the cholinergic MGINs.The MGINs then activate the cholinergic motor neurons to evoke swimming (Figure 1A and B). The significance of the non-AMPAR glutamatergic input to the pr-AMG RNs is explored in the present study.

The circuit logic for the PR-II mediated dimming response is more complex. GABAergic projections from the PR-IIs are targeted exclusively to the GABAergic pr-AMG RNs (Figures 1B and C). This arrangement led to a disinhibitory model for the light-dimming response (Kourakis et al., 2019). In this model, swimming is actively inhibited by pr-AMG RN input to the MGINs unless they are themselves inhibited by the GABAergic photoreceptors (Figure 1B). Tunicate photoreceptors, like their vertebrate counterparts, are hyperpolarizing (Gorman et al., 1971), and thus dimming would be expected to increase their GABA release. Moreover, behavioral analyses with the GABA antagonist picrotoxin, as well as in the mutant *frimousse*, in which aBV is transfated to epidermis (Hackley et al., 2013), indicate that swimming behavior is constitutively inhibited, consistent with the disinhibition model.

Both *Ciona* visuomotor behaviors are responses to changing illuminations, whether it be decreased ambient illumination for the PR-II circuit, or directional photoreceptor shading in the PR-I circuit. For both visuomotor circuits to function in changing illumination conditions, dynamic visual processing is required. We report here that the *Ciona* larval CNS processes visual inputs to detect fold-change differences. In fold-change detection (FCD), the response depends only on the relative change in input, and not on the absolute change (Adler and Alon, 2018; Alon, 2019). FCD allows a sensory system to give a consistent response to the same relative change, independent of the ambient conditions, while suppressing noise. Moreover, we present evidence that the circuits for FCD are distinct from the adaptive mechanisms of the photoreceptors, and instead appear to be present in the complex of synaptic connectivity between relay neurons in the pBV. Finally, we note that the convergence of conserved gene expression, anatomical location, and most significantly, function and connectivity, all point to the pBV sharing deep homology with the vertebrate midbrain visual processing centers, including the tectum and pretectum. While gene expression patterns have been presented as evidence of a tunicate midbrain homolog (Imai et al., 2002, 2009), it has been more widely speculated that tunicates lack a midbrain homolog (Cañestro et al., 2005; Holland, 2009; Ikuta and Saiga, 2007; Lacalli, 2006; Satoh and Levine, 2005; Striedter and Northcutt, 2020; Takahashi and Holland, 2004).

## Results

### Larval visuomotor behaviors display fold-change detection

To assess *Ciona* larval visuomotor behaviors, 25-hour post-fertilization larvae were placed in 6 cm petri dishes (up to 300 larvae at one time) and illuminated with a programmable 505 nm LED lamp at varying intensities while recording at far-red (700 nm), as described previously (Bostwick et al., 2020; Kourakis et al., 2019). In the first set of experiments, the response of larvae to a light-dimming series from 3-fold (600 lux to 200 lux) to 60-fold (600 lux to 10 lux) was assessed. Several parameters of the dimming-induced swims were measured: the percent of larvae responding to dim and their reaction time, as well as the duration, speed, and tortuosity of swims. Movie 1 shows representative responses to 3-, 10− and 60-fold dims. Of these parameters, induced swim duration showed a positive relation to increased fold-change (Figure 2A), while speed and tortuosity were constant across the series (Figure 2B and C). The percent of larvae responding to dimming also did not track with the fold-change series. The percent responding increased initially at the lowest fold-changes, but plateaued at around 10-fold with ~100% of larvae responding (Figure 2D).

**Figure 2.**
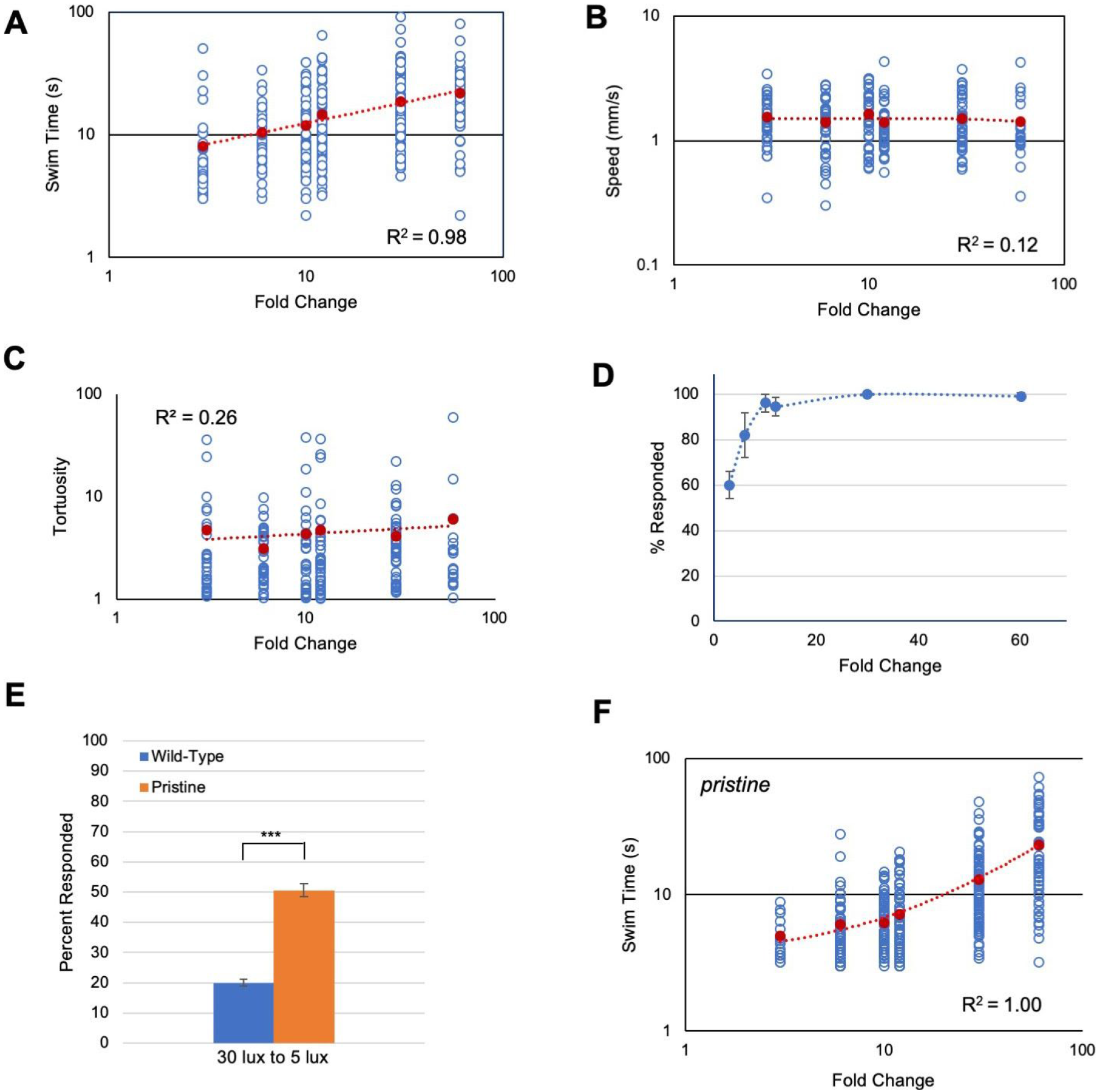
Response of *Ciona* larvae to fold-change light dims. **A.** Larval swim times increase as a power function in response to increased fold-change light dimming (3-fold to 60-fold; log/log plot shown). All data points (blue circles) and averages (red circles) are shown (same for panels B, C, and F). See **Movie 1** for representative results. **B**. Swim speed is constant across fold-change dimming series. **C.** Swim tortuosity is constant across fold-change dimming series. **D.** The percent of larvae responding as a function of fold-change dimming. Shown in graph are the averages from three recordings (± S.D.). **E.** Percent of larvae responding to six-fold dim at low illumination conditions (30 lux to 5 lux). *n*= 76 and 131, for wild-type and *pristine*, respectively. **F.** Larval swim times of homozygous *pristine* mutants increase as a linear function in response to increased fold-change light dimming (3-fold to 60-fold; log/log plot shown). (*** = p < 0.001) See **Figure 2 - Source Data 1. Fold Change Detection Behavior** for sample sizes, average values, and statistical analyses.

While the dimming response is an output of the PR-II circuit, the PR-I negative phototaxis circuit depends on the larvae detecting changing illumination as they perform casting swims. Although we have reported that *Ciona* larvae are able to successfully navigate in a wide range of ambient lighting conditions (Salas et al., 2018), the phototaxis assay would not permit precise control of the amount of light the PR-I photoreceptors were receiving, making it difficult to assess their responses to fold-change stimuli. To circumvent this problem, we used the loss-of-pigmentation mutant *pristine* (*prs*) to assess the PR-I photoreceptors, as we have done previously (Kourakis et al., 2019; Salas et al., 2018). In larvae homozygous for *prs* the PR-I photoreceptors respond to ambient light changes because they are no longer shielded by the pigment cell; in other words, changes in ambient light mimic casting swims (Salas et al., 2018). While both the PR-I and PR-II photoreceptors would be stimulated by dimming in *prs* mutants, there are more Group I photoreceptors than Group II (23 versus 7), and the Group I output appears to predominate (Salas et al., 2018). One way this is evident is that the dimming-evoked swims of *prs* mutants are straight, as are phototaxis swims, rather than highly tortuous, as are dimming-induced swims (Salas et al., 2018). To validate this further, we find that *prs* larvae are more sensitive to dimming at low-light conditions than wild type larvae (Figure 2E), consistent with the behavioral output from *prs* mutants primarily reflecting the output from the PR-I circuit.

When the fold-change dimming series was performed on *prs* larvae, we again observed a positive relationship of swim time to fold-change, but with a significantly different shape to the response curve (Figure 2F and Figure 2 - Source Data). Modeling indicates that a number of different circuit motifs, including the incoherent type-1 feedforward loop (I1FFL) and the nonlinear integral feedback loop (NLIFL), can generate FCD outputs (Adler et al., 2017). Moreover, different FCD circuit motifs can generate different response curves (*e.g.*, linear or power), meaning that the response curves can be diagnostic of the underlying circuit architecture (Adler and Alon, 2018). For wild type *Ciona* larvae, the curve of swim time versus fold-change was found to best fit a power function, with R^2^ = 0.98 (Figure 2A). While a log function also fit this curve with R^2^ = 0.98, the Bayesian information criterion (BIC, see Methods) for a power relationship had the lower score, indicating a better fit (−13 and 17 for power and log, respectively; R^2^ = 0.87 for linear). The best fitting model for the *prs* swim time responses was a linear curve having an R^2^ value equal to 1.00 (R^2^ = 0.92 and 0.81 for power and log, respectively). In summary, wild-type and *prs* larvae both show a positive relationship between fold-change dimming and swim time, although with different response curves, suggesting that different FCD circuits may be responsible.

### Validation of FCD behavior

FCD mechanisms, while incorporating widely observed phenomena such as adaptation and log transformation, have distinct attributes, the most important being scale-invariance (Goentoro et al., 2009; Kamino et al., 2017; Shoval et al., 2010). With scale-invariance the output depends only on the fold-change, not on the absolute magnitude of the stimulus. We find scale-invariance holds true for the *Ciona* visuomotor response across at least three orders of magnitude. To assess scale-invariance, wild-type and *prs* larvae were exposed to series of 3-, 10− and 60-fold dims, but from starting intensities of 3000, 300 and 30 lux (*e.g.*, the 10-fold dims were 3000 to 300 lux, 300 to 30 lux, and 30 to 3 lux). We observed that the swim time responses of both wild-type and *prs* larvae were not significantly different within a fold-change, irrespective of the magnitude of illumination, but were significantly increased as fold-change increased (Figure 3A and B and Figure 3 - Source Data).

**Figure 3.**
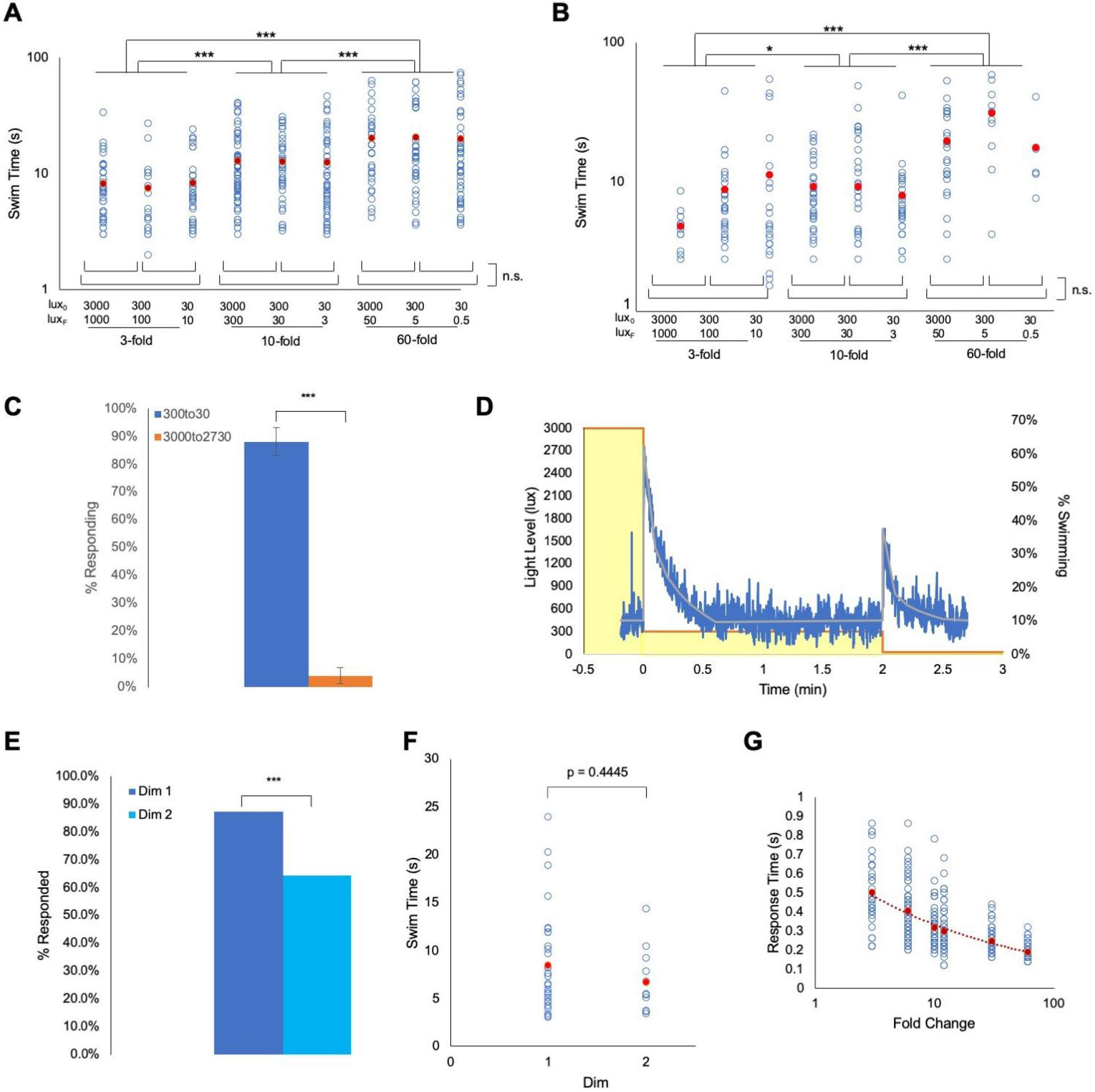
*Ciona* fold-change detection response. **A.** *Ciona* larvae swim times show scale-invariance to light dimming across three orders of magnitude. Lux_0_: initial illumination level in lux; Lux_F_: illumination after dim in lux. **B.** Same light-dimming series as in panel A, but with *pristine* mutants. **C.** The *Ciona* light-dimming response follows Weber’s law. Shown are the percent of larvae responding to 270 lux dimming from the initial conditions of 300 lux or 3000 lux (n = 776 and 443; respectively). **D.** *Ciona* larvae show absolute adaptation to light-dimming. Larvae were exposed to two light dimmings separated by two minutes (3000 lux to 300 lux, and 300 lux to 30 lux; yellow boxes). The blue line shows the percent of larvae swimming at 1 second time points (n = 91-320), and the grey line is the average at each time point. See also **Movie 2. E.** For data shown in panel D, a higher percentage of larvae responded to the 3000 to 300 lux dim (Dim 1) than to the 300 to 30 lux dim (Dim 2). **F.** For the experiment shown in panel D, the swim times induced by the 3000 to 300 lux dim (Dim 1) were not different from the 300 to 30 lux dim (Dim 2). **G.** Plot of the reaction time versus fold change. This is defined as the time point at which swimming was first detected following dimming. For panels A, B, F and G, all data points (blue circles) and averages (red circles) are shown. See **Figure 3 - Source Data 1. Validation of fold change detection behavior** for full data and statistical analyses. (*** = p < 0.001; * = p < 0.05; n.s. = not significant).

Integral to fold-change detection behavior is Weber’s law (*i.e.*, the change in stimulus needed to elicit a response is proportional to the absolute value of the original stimulus) (Adler and Alon, 2018). To demonstrate this directly in *Ciona* visuomotor behavior, larvae were adapted to either 3000 lux or 300 lux were then dimmed by 270 lux (*i.e.*, to 2730 or 30 lux, respectively). In those larvae adapted to 3000 lux, we observed no response to the dimming, while in the larvae adapted to 300 lux, we observed vigorous swimming in nearly all larvae (Figure 3C). Another property of FCD systems is exact adaptation [*i.e.*, the system returns to the baseline state even when the modulated stimulus persists at the new state (Shoval et al., 2010)]. To investigate this, larvae were initially adapted to 3000 lux, which was then dimmed to 300 lux and held at this level for 2 minutes. The illumination was then dimmed a second time, to 30 lux (Movie 2). Figure 3D shows a plot of swimming activity of the larvae as illumination levels change. We observed that the larvae respond robustly to the first 10-fold dim (3000 to 300 lux, yellow boxes in Figure 3D), but stop swimming after approximately 30 seconds. The majority of the larvae were then stationary until the second 10-fold dimming (300 to 30 lux). While a lower percent of larvae responded to the second dim than the first (87% vs 64%, Figure 3E), no difference in the average swim times of responding larvae was observed (Figure 3F). Finally, another predicted property of FCD systems is that the reaction time should be inversely proportional to the fold change (Adler and Alon, 2018), which we observed as a power-slope increase in reaction time as the fold-change decreased (Figure 3G and Figure 3 - Source Data).

### Fold-change detection circuits in the Ciona *CNS*

FCD mechanisms have been primarily studied in the context of signal transduction and microbial chemotaxis, rather than in animal sensory systems (Adler and Alon, 2018). The clearest case for apparent FCD in an animal sensory system is in *C. elegans* chemotaxis, where the fold-change transformation of sensory input appears to be a property of the chemoreceptors, although the precise intracellular circuit is not known (Larsch et al., 2015). For visuomotor systems, such as those characterized here for *Ciona* larvae, the most obvious location for the FCD circuit would be the photoreceptors. However, while vertebrate photoreceptors (particularly cones) obey Weber’s law (Burkhardt, 1994; Fain et al., 2001), FCD by phototransduction machinery has not been reported. Moreover, modeling suggests that adaptation and adherence to Weber’s law alone are not sufficient to give FCD (Shoval et al., 2010). While the calcium-dependent mechanism of photoreceptor adaptation is well characterized (Pugh and Lamb, 1990), results presented here suggest that more than one qualitatively different FCD circuit may be in operation in the *Ciona* visual systems (Figure 2). Finally, the adaptive properties of the vertebrate visual system arises not only from the transduction mechanism inherent to the photoreceptors, but also from the neural circuitry in the vertebrate retina (Dunn et al., 2007). By contrast, the *Ciona* ocellus lacks the complex interneuron architecture of the vertebrate retina, rather the photoreceptors project directly to the pBV. Taken together, these observations suggest that a wider consideration of the *Ciona* CNS, beyond the photoreceptors, may yield new insight into FCD mechanisms.

The pBV (Figure 1A) is a likely location for components of the FCD circuits. The pBV receives input not only from the two photoreceptor systems, but also from the gravity-sensitive otolith organ via projections from the glutamatergic antenna cells, as well as from other sensory systems (*e.g.*, peripheral nerves and coronet cells) (Ryan et al., 2016). The synaptic connectivity between the *antenna cell relay neurons* and prAMG RNs in the pBV appears to integrate visual and gravitaxis inputs, resulting in a behavior in which negative gravitaxis (*i.e.*, upward swimming) is triggered by light dimming (Bostwick et al., 2020). Thus, the pBV appears to function not only as a simple relay for sensory information, but also a sensory processing center. Figure 4A shows the full PR-I and PR-II visuomotor circuits, as given by the *Ciona* connectome (Ryan et al., 2016) with superimposed putative neurotransmitter types, as deduced by *in situ* hybridization (Kourakis et al., 2019). When the PR-I and PR-II circuits are simplified by combining cell types and synaptic connections, two plausible FCD circuits are evident, with prominent roles played by the pBV relay neurons (Figure 4B). The proposed PR-I circuit contains an apparent incoherent feedforward loop, but differs from an I1FFL in having an additional excitatory interaction from the output (y) to the modulatory (m) (top, Figure 4B). Computer modeling of circuit motifs indicates that this configuration should give FCD (circuit 04040 in Adler et al., 2017). While the I1FFL motif should result in a log function for input-output relationship (*e.g.*, fold-light dimming and swim time) (Adler et al., 2014), our observations for *prs* larvae best fit a linear curve (Figure 1F). It is not known if the proposed circuit motif for the PR-I circuit would predict a linear relationship, but it is clear that it does not conform to the I1FFL prediction, and that, most significantly, the relationship differs from observations for the PR-II circuit. The PR-II circuit, being disinhibitory, is more unusual, but resembles a NLIFL, but with the modulator (m) being excitatory rather than inhibitory (bottom, Figure 4B). Moreover, the input-output relationship (*e.g.*, fold-light dimming and swim time) for circuits containing the NLIFL motif follow a power relationship (Adler et al., 2014), as was observed (Figure 1A). Finally, we note that in the two proposed FCD circuits in Figure 4B that the RN types exchange roles as output (y) and modulator, suggesting an economy of neuron use.

**Figure 4.**
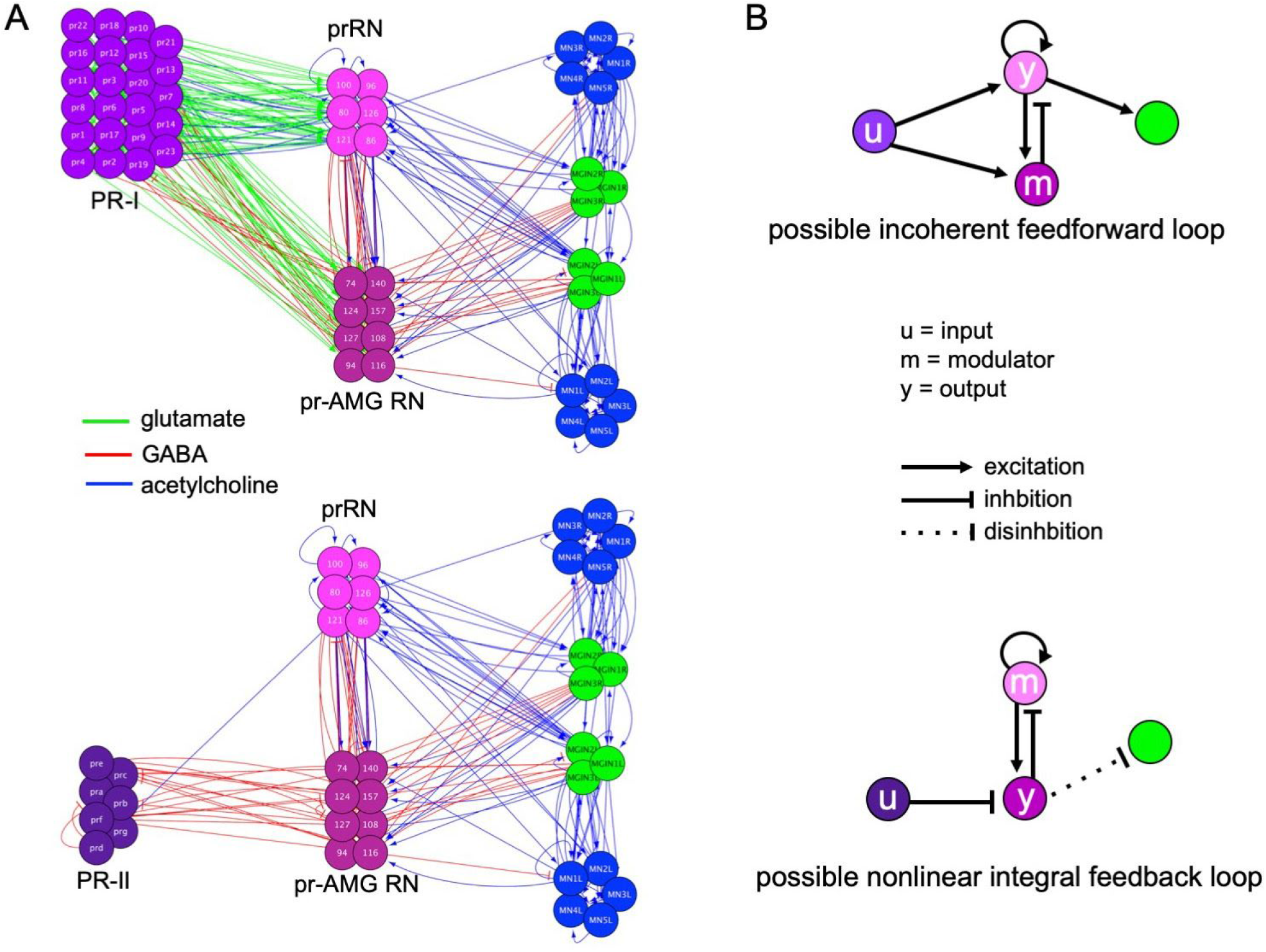
Visuomotor circuits and putative fold-change detection circuits. **A.** Full circuits for the PR-I (top) and PR-II (bottom) pathways from the *Ciona* connectome (Ryan et al., 2016). Neurotransmitter use for synaptic connections (lines) is based on Kourakis et al., 2019. Abbreviations: PR, photoreceptor; prRN, photoreceptor relay neuron; pr-AMG RN, photoreceptor ascending motor ganglion relay neurons; MGIN, motor ganglion interneurons; MN, motor neurons. **B.** Simplified circuits for the PR-I (top) and PR-II (bottom) pathways derived by combining like cells and assigning valence of synapses (excitatory or inhibitory) based on consensus for that cell type. Nodes are labeled according to proposed function (*i.e.*, input modulator, and output). Colors of neuron classes are according to (Ryan and Meinertzhagen, 2019).

In the proposed PR-II NLIFL FCD circuit the modulators are the cholinergic prRNs (Figure 4B). Multiplex *in situ* hybridization showed that the AMPAR is coexpressed with VACHT in the prRNs, but not in the VGAT-positive pr-AMG RNs (Kourakis et al., 2019). Accordingly, treatment of larvae with the AMPAR antagonist perampanel inhibits the PR-I mediated phototaxis, but not the PR-II mediated dimming response (Kourakis et al., 2019). We report here that AMPAR is not expressed in the photoreceptors, but is expressed in the motor ganglion as well as in the pBV (Figure 5A). In the proposed NLIFL circuit the output cells (the GABAergic pr-AMG RNs) inhibit the cholinergic prRN modulators (Figure 4). We hypothesized that by adding AMPA, an AMPAR agonist, the activity of the modulator, and therefore FCD, should be disrupted, but not the ability of the larvae to respond to dimming. Our results agreed with the hypothesis. We observed no difference in the percentage of control and AMPA-treated (500 μm) larvae responding to a fold-dimming series (Figure 5B). However, a plot of swim time versus fold change (Figure 5C) shows the slopes of the two curves were significantly different (R^2^ of 0.98 for control, and 0.75 for AMPA-treated). In a second set of experiments in which control and AMPA-treated larvae were assessed against a series of identical fold-changes but of different magnitudes, the disruption to the FCD mechanism was evident (Figure 5D and E). When the data were grouped according to fold-change, no significant differences in swim times were found between fold changes (Figure 5D), unlike in untreated larvae (Figure 3A). However, when the data were grouped by magnitude of the starting illumination, the larvae appeared to be responding according to the magnitude of illumination, rather than fold change (Figure 5E). For example, larvae assessed from a starting illumination of 30 lux had shorter swims than larvae assessed from a starting illumination of 300 lux, independent of the fold change. This was true for comparisons across all starting illuminations, with the exception of 300 lux versus 3000 lux.

**Figure 5.**
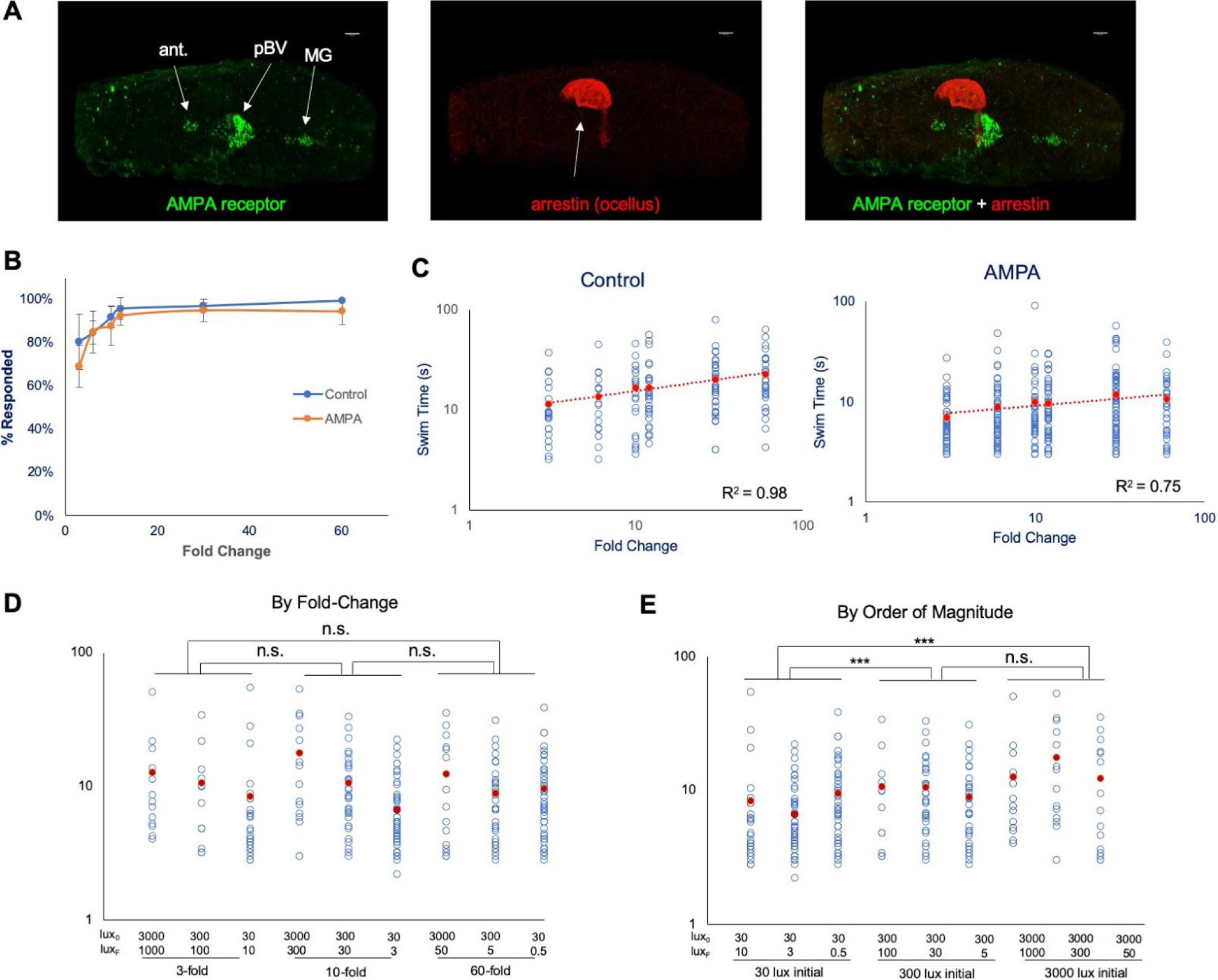
Disruption of FCD by an AMPA receptor agonist. **A.** AMPA receptor expression detected by *in situ* hybridization (green), and the ocellus photoreceptor marker Arrestin was detected by immunostaining (red). Dorsal view, anterior to left. Abbreviations: ant., antenna cells; pBV, posterior brain vesicle; MG, motor ganglion. **B.** Percentage of control and AMPA-treated larvae responding to the indicated fold-change dimmings. The averages from three recordings are shown (± S.D.). **C** Swim times of control and AMPA-treated larvae in the fold-change dimming series. The R^2^ valves for the curves are indicated. **D.** AMPA-treated larvae do not show scale-invariance to fold-change dims of different magnitudes (compare to controls in Figure 3A). **E.** Data from panel D sorted by magnitude of initial illumination (lux_0_). (*** = p < 0.001; n.s. = not significant). For panels C, D, E all data points (blue circles) and averages (red circles) are shown. See **Figure 5 - Source Data 1 AMPA disrupts fold change detection** for full data and statistical analyses.

In summary, these results show that AMPA treatment can disrupt the FCD mechanism, without changing the ability of the larvae to respond to dimming. Moreover, the absence of AMPARs in the photoreceptors indicates that at least some components of the FCD machinery lie outside of the photoreceptors. We favor the pBV as the likely site for the FCD circuits for two reasons. First, published results already point to the pBV as a sensory processing center (Bostwick et al., 2020). Second, not only have we identified putative FCD circuits in the pBV, but the architecture of the proposed PR-I and PR-II circuits are different, as predicted by the FCD response curves (Figure 1). Nevertheless, we also observed AMPA receptors in the MG, and we can not exclude a contribution from neurons there in the observed AMPA treatment results. However, analysis of the connectome in MG did not reveal any plausible FCD circuits. The MG is dominated by excitatory cholinergic interneurons and motor neurons, while inhibitory neurons, which would be an essential modulatory element of any likely FCD circuit, are limited to the GABAergic AMG neurons, which received no descending input, directly or indirectly, from the photoreceptors or the BV, and the glycinergic decussating ACINs, which likely play a role in the central pattern generator, not visual processing (Kourakis et al., 2019; Nishino et al., 2010; Ryan et al., 2016). In addition, the PR-I and PR-II circuits project from the pBV on a common set of MG interneurons, making it unlikely that this brain region would be responsible for the different fold-change response curves for the PR-I and PR-II circuits.

### Is the pBV a homolog of the vertebrate midbrain?

The pBV is the primary recipient of projections from the ocellus, otolith, and coronet cells, and a subset of peripheral neurons (Ryan et al., 2016). Relay neurons within the pBV then project posteriorly through the neck to the MG. No other region of the *Ciona* CNS has this convergence of sensory inputs and descending interneuron projections. Results presented here, as well as published studies (Bostwick et al., 2020), point to the pBV as a sensory processing and integrating center. Thus in many ways the function of the pBV resembles that of the vertebrate midbrain visual processing centers, including the optic tectum (Knudsen, 2020). The resemblance of the pBV to the vertebrate midbrain extends to the *Ciona* CNS anatomy as well. In particular, the pBV is located immediately anterior to the neck region, which based on gene expression, and the fact that it forms a constriction in the CNS, is thought to have homology to the vertebrate midbrain-hindbrain junction (Ikuta and Saiga, 2007). Despite these anatomical similarities, it is widely speculated that tunicates either do not have, or have lost a midbrain homolog. These reports are based on the expression patterns of several genes that do not match those of vertebrates. For example, the gene DMBX, which plays an essential role in vertebrate midbrain development, is not expressed anterior to the MG in *Ciona* (Ikuta and Saiga, 2007; Takahashi and Holland, 2004). In addition, the tunicate *Oikopleura dioica* (Class Larvacea) does not express the genes *engrailed* or *pax2/5/8* anterior to its hindbrain, suggesting that larvaceans lack a midbrain (Cañestro et al., 2005). However, these studies were limited to a few genes, and were performed before the connectivity of the pBV was made apparent by the publication of the connectome. Moreover as presented below, a wider view of neural genes shows extensive gene expression conservation between the pBV and the vertebrate midbrain.

The *Ciona* BV is divided into distinct anterior and posterior domains that derive from invariant cell lineages arising at the 8-cell stage, with the anterior BV (aBV) descending from the a-lineage, and the pBV from the A-lineage (Figure 6A, red and blue cell centroids, respectively) (Nicol and Meinertzhagen, 1988; Nishida, 1987). Moreover, the distribution of neuron types is sharply demarcated by this boundary, with the relay neurons, which uniquely project from the BV to the MG, being found only in the pBV. The relay neurons are themselves segregated within the pBV, with those receiving photoreceptor input clustered anteriorly (Figure 6B). The gene *Otx*, which is expressed in the forebrain and midbrain of vertebrates (Boyl et al., 2001), is expressed in *Ciona* in both the aBV and pBV (Hudson et al., 2003; Ikuta and Saiga, 2007; Imai et al., 2002), while a number of the vertebrate forebrain markers are expressed only in the aBV lineage. This includes the genes *Dmrt1* (Kikkawa et al., 2013; Tresser et al., 2010; Wagner and Levine, 2012), as well as *Lhx5*, *Six3*, and *Gsx2* (Esposito et al., 2017; Mazet et al., 2005; Moret et al., 2005; Reeves et al., 2017), all of which play essential roles in vertebrate forebrain development (Kirkeby et al., 2012; Lagutin, 2003; Peng, 2006; Toresson et al., 2000). While *Gsx2* is also expressed in the pBV, this is only in later stages of development (tailbud stages). Additional vertebrate forebrain markers expressed exclusively in the aBV lineage include *Lhx2/9*, *Bsx* (or *Bsh*) and *Arx* (or *Aristaless*) (Cao et al., 2019). In vertebrates these genes are reported to play essential roles in cortex development (Marsh et al., 2016; Roy et al., 2014; Schredelseker et al., 2020; Shetty et al., 2013).

**Figure 6.**
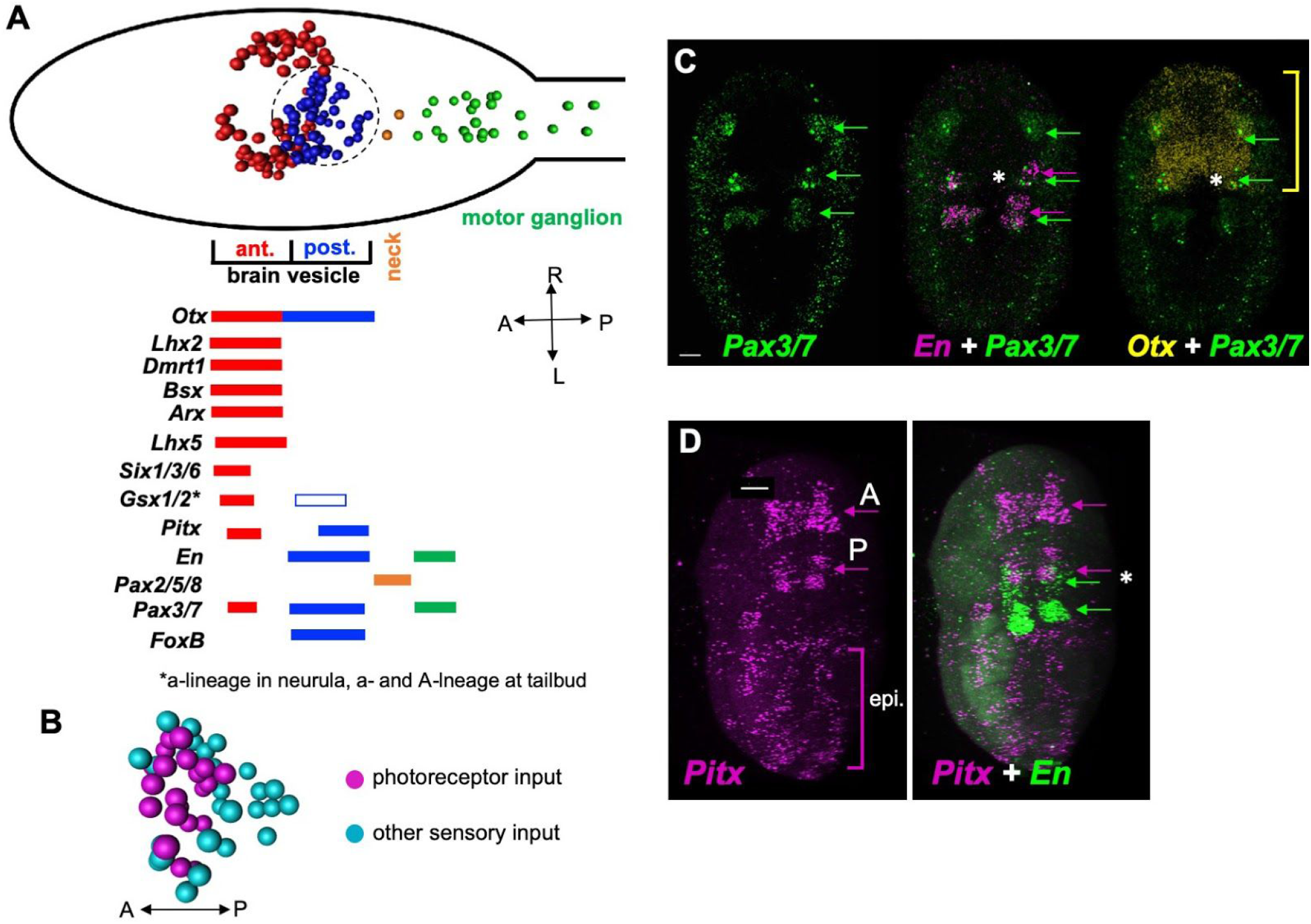
Gene expression in the *Ciona* posterior brain vesicle suggests homology with the vertebrate midbrain. **A.** Top: Diagram of the *Ciona* larval CNS with the major brain regions indicated in color. The centroids of neurons are shown (from Ryan et al., 2016). Bottom: summary of embryonic gene expression patterns marked by corresponding larval CNS domains. **B.** Spatial distribution of relay neuron types in the pBV. Region shown corresponds to the circled area in panel A (centroids shown). **C.** *In situ* hybridization for *Pax3/7*, *En*, and *Otx* in early tailbud *Ciona* embryos. The anterior domain of *En*, which marks the presumptive pBV, overlaps with *Pax3/7* (asterisk middle panel), and *En* (asterisk right panel). Yellow bracket shows anterior-posterior extent of Otx expression. **D.** *In situ* hybridization for *Pitx* and *En*. *Pitx* is expressed in anterior and posterior domains in early tailbud embryos. The posterior domain of *Pitx* overlaps with the anterior *En* domain in the pBV (asterisk right panel). Consistent with previous reports, labeling was also observed in the epidermis. Abbreviations: A, anterior; P, posterior; epi, epidermis. Anterior is to the left for panel A and D, and to the top for B and C. Scale bars are 10 microns.

By contrast, the pBV expresses a number of genes associated with the vertebrate midbrain. This includes the A-lineage specific marker *FoxB1* (or *Ci-FoxB*) (Moret et al., 2005; Oonuma et al., 2016), which in vertebrates plays a role in midbrain development (Wehr et al., 1997). The development of the vertebrate optic tectum, a midbrain structure, requires the co-expression of *Pax3*, *Pax7*, *Otx2*, and *En* (Matsunaga et al., 2001; Thompson et al., 2008). The *Ciona engrailed* homolog is expressed in two domains embryologically, posteriorly in the MG and anteriorly in the pBV (Ikuta and Saiga, 2007; Imai et al., 2002). We observed overlapping expression of *Pax3/7*, *En*, and *Otx2* in the developing pBV of early tailbud embryos (Figure 6C, asterisk). Finally, *Pitx* has a well-defined role in vertebrate midbrain development (Luk et al., 2013). We observed *Ciona pitx* expression at early tailbud stage in a posterior domain that overlaps with pBV *engrailed* expression (Figure 6D, asterisk), as well as in an anterior domain that appears to correspond to the aBV expression reported in older embryos and larvae (Christiaen et al., 2002). In addition, diffuse epidermal labeling was observed, as reported previously (Boorman and Shimeld, 2002). As reported previously, the pBV is bounded posteriorly by expression of *Pax2/5/8* in the neck cells, indicating homology with the vertebrate MHB (Ikuta and Saiga, 2007) (Figure 6A).

These expression pattern results show that the BV has distinct anterior and posterior expression domains, with the anterior domain expressing genes known to be expressed in the developing vertebrate forebrain, and the posterior domain expressing genes associated with the developing vertebrate midbrain. These observations do not agree with previous reports that suggest the entire BV is homologous to the vertebrate forebrain (and that a midbrain homolog is absent). Thus the convergence of gene expression, anatomical, connectivity and functional data all point to the pBV as sharing a common origin with the vertebrate midbrain. Reaching deeper into chordate history, evidence has been presented that cephalochordates, the third chordate subphylum which diverged before the split of the tunicates and vertebrates, have conserved midbrain visual processing centers (Lacalli, 2006; Suzuki et al., 2015; Takahashi and Holland, 2004). Thus the scenario by which tunicates lack a midbrain homolog would appear to require them losing this structure, and then evolving *de novo* a brain structure in a convergent location with convergent connectivity.

## Discussion

Behavioral studies presented here show that *Ciona* larvae process visual input to detect fold-change. The utility of this behavior is clear: in negative phototaxis, larvae discern the direction of light via casting swims, and it is the change in illumination falling on the PR-Is as the larvae turn away from the light that is the cue to swim. FCD ensures that the casting mechanism functions in the wide range of ambient light conditions that the larva is likely to encounter, and that the response is invariant to the scale of the input. The function of FCD in the light-dimming response is similar. In the absence of FCD, the change in illumination caused by the same looming object might appear as a threat in one ambient light condition, but not under different lighting conditions. FCD ensures that the response varies as a function of the relative shading caused by the looming object.

A number of cellular signaling systems have been shown to give FCD responses to extracellular cues, including those in bacterial chemotaxis and growth factor signaling in mammalian cells and embryos (Adler and Alon, 2018; Goentoro et al., 2009; Lyashenko et al., 2020; Shoval et al., 2010), and modeling has identified several classes of biological circuits that can generate FCD responses (Adler et al., 2017; Goentoro et al., 2009; Hironaka and Morishita, 2014). Here we identified components of the *Ciona* CNS that appear to be responsible for the FCD processing. The *Ciona* photoreceptors themselves, through their adaptive mechanisms, certainly play a role in processing the visual inputs. However, we present evidence that neural circuits, and not just signal transduction pathways, can generate FCD outputs. Incoherent feedforward motifs, including I1FFL motifs, are common in neural circuits, although they have not been assessed for FCD (Cognigni et al., 2018; Feldmeyer et al., 2018; Ishimoto and Kamikouchi, 2020; Jeanne and Wilson, 2015). As evidence for neural circuitry generating a FCD response, we report here that treatment of *Ciona* larvae with the AMPAR agonist AMPA was able to transform scale-independent behavior of the dimming response to one that scaled with the magnitude of the illumination. Based on the behavior of the AMPA-treated larvae, we hypothesize that AMPA targets the inhibitory modulator (m) of the circuit, which acts as a repressor of the output neurons (y), but not the output neurons themselves (Figure 4B). The distribution of AMPARs and of plausible FCD circuits points to the pBV as a major contributor to FCD.

The conservation of function, connectivity, anatomy and gene expression are consistent with the pBV having a common origin with the visual process centers in the vertebrate midbrain. Moreover, evidence has been presented that cephalochordates (*e.g.*, amphioxus), which comprise a third chordate subphylum that diverged before the split of the tunicates and vertebrates, also have a brain structure which shares a common origin and function with the vertebrate midbrain (Albuixech-Crespo et al., 2017; Holland, 2015; Lacalli, 2018). If this hypothesis is borne out, the scenario by which tunicates would lack a midbrain homolog would require a loss of this structure, followed by *de novo* evolution of a sensory processing center in a convergent location with convergent connectivity, and with largely convergent gene expression. A simpler explanation is that the origins of the midbrain go back to the base of the chordates. Thus, observations of *Ciona* behavior and neural connectivity may provide insight into ancestral and conserved visual processing mechanisms. Shared midbrain homology across chordates, and additional evidence of still deeper brain homology, including possible tripartite organization and an MHB precursor in the protostome-deuterostome ancestor (Hirth et al., 2003; Bridi et al., 2020), suggest that vertebrate CNS patterning and overall regional architecture may trace more ancient roots than has generally been appreciated.

## Method

### Animals

Wild type *Ciona robusta* (a.k.a., *Ciona intestinalis* type A) were collected from the Santa Barbara harbor. The animals carrying the mutation *pristine* (Salas et al., 2018) were cultured at the UC Santa Barbara Marine Lab (Veeman et al., 2011). Larvae were obtained by mixing dissected gametes of 3 adults and then culturing in natural seawater at 18°C. Homozygous *prs* larvae were produced by natural spawning of heterozygous or homozygous *prs* adults. For Fig. 1C, two stable transgenic lines, vgat>kaede and vacht>CFP [provided by Y. Sasakura], were crossed to yield offspring with labeled GABAergic/glycinergic cells and cholinergic cells, respectively.

### Hybridization chain reaction (HCR) in situ and immunolabeling

Whole mount *in situ* hybridization of embryonic or larval *C. robusta* were performed as previously described (Kourakis et al., 2019). Briefly, fluorescent labeling was achieved following the hybridization chain reaction (v. 3.0) of Molecular Instruments (Los Angeles).

Complementary RNA probe sets were designed to coding regions for the following *C. robusta* genes: *Otx*, *en*, *pax3/7*, *AMPA receptor*, and *pitx*. In larvae which underwent both *in situ* labeling and immunostaining, the *in situ* hybridization was performed first, followed by the immunolabeling (see below), after a transition from 5X SSCT to PBST.

Larvae for immunostaining were dechorionated at mid-tailbud stage using sodium thioglycolate, as for *in situ* hybridization, so that left-right asymmetric properties of the CNS would not be disrupted (Yoshida and Saiga, 2008). The immunostaining followed previously described procedures for *Ciona* (Newman-Smith et al., 2015). A primary antibody against *C. robusta* arrestin (Horie et al., 2005), raised in rabbit, was used at a dilution of 1:1000. Α secondary antibody, α-rabbit AlexaFluor 594 (Invitrogen), was also used at 1:1000. For vgat>kaede and vacht>CFP larvae, rabbit α-Kaede (MBL) and mouse α-GFP (Life Technologies) antibodies were used at 1:1000, followed with appropriate AlexaFluor secondaries, also at 1:1000 dilution (described above).

Labeled animals (either by *in situ* or immunohistochemistry) were imaged on an Olympus Fluoview 1000 confocal microscope; post-image analysis used Imaris v6.4.0.0 or ImarisViewer v9.5.1 as well as Fiji (ImageJ) v. 2.0.0-rc-69/1.52p. The surface model depicted in Fig. 1C was generated in Imaris v6.4.0.

### Behavioral Assays

All larvae were between 25 and 28 hours post fertilization (hpf) (18°C). Larval swimming behaviors were recorded in sea water with 0.1% BSA using agarose-coated petri dishes to reduce sticking. Image series were collected using a Hamamatsu Orca-ER camera fitted on a Navitar 7000 macro zoom lens. Programmable 700 nm and 505 nm LED lamps (Mightex) mounted above the petri dishes were used for dimming response assays as previously described (Kourakis et al., 2019). The dim response, adaptation, and reaction time movies were recorded at 5, 8.9, and 50 frames per second (fps), respectively. In the standard assay larvae were recorded for 10s at the initial intensity (lux0) that was then dimmed (luxF) to specific values while image capture continued for 2 min. Larvae were allowed to recover for 5 min before being assayed again. All light intensity readings were taken with an Extech Instruments light meter.

### Drug Treatments

(*RS*)-AMPA hydrobromide (Tocris) was dissolved in filtered sea water to a stock concentration of 7.5 mM and then diluted to a final concentration of 500 µM. Larvae were incubated with the drug for about 10 min before beginning assays.

### Behavioral Quantification

Larvae with short bouts of swimming (<3s) were not scored (Bostwick et al., 2020; Kourakis et al., 2019). Swim times, speeds, and tortuosities were calculated using the MATLAB script ELIANE (Kourakis et al., 2019) and the larvae scored between the conditions were tested for significance using the Wilcoxon Rank-Sum test. In the test for absolute adaptation (Figure 2), to measure the percent of larvae swimming at 1 second time intervals, the ELIANE script was modified to determine if a centroid (*i.e.*, larva) in frame x was in the same position in frame x + 1. The sum of moving centroids was then divided by the total number of centroids. Percent of larvae that responded to dimming stimuli was quantified manually and tested for significance using the Student T-Test. R-squared values and Bayesian information criterion (BIC) were calculated using the program R. Larvae that had stuck to the petri dish, or that swam to the edges or into other larvae were not scored.

## Acknowledgments

The work was supported by an award from the NIH (NS103774) to W.C.S, and a CCS Create Fund Summer Fellowship to S.S.. We thank Takehiro Kusakabe for the anti-Arrestin antibody.

## Author contributions

C.B. and M.J.K. contributed to experimental design, data collection and analysis, and manuscript preparation. S.S. and L.B. contributed to data collection and analysis. W.C.S. contributed to research funding, experimental design, data collection and analysis, and manuscript preparation.

---

**Movie 1. Response of *Ciona* larvae to 3-, 10− and 60-fold dimminge**. Larvae in 60 mm petri dishes are recorded from above. All larvae were initially exposed to a 505 nm LED lamp at 600 lux. The lamp was then dimmed to 200, 60 or 10 lux, as indicated. The swimming behavior of 10 larvae from each group is tracked (yellow boxes). The swim track is projected across time in color, and the yellow boxes disappear when the larvae stop swimming. Notice that the swims are progressively longer as larvae exposed to the larger fold changes (*i.e.*, the projected swims in color are longer). The larvae were recorded using a far red light (700 nm). Movie plays at normal speed.

**Movie 2. Absolute adaptation.** Larvae are exposed to two dimming events separated by 2 minutes. The first dim is from 3000 to 300 lux, and the second from 300 lux to 30. Notice that swims are induced at each dimming, but that the swimming largely stops between dims. Video plays at 5X normal speed.

**Figure 2 - Source Data 1.**
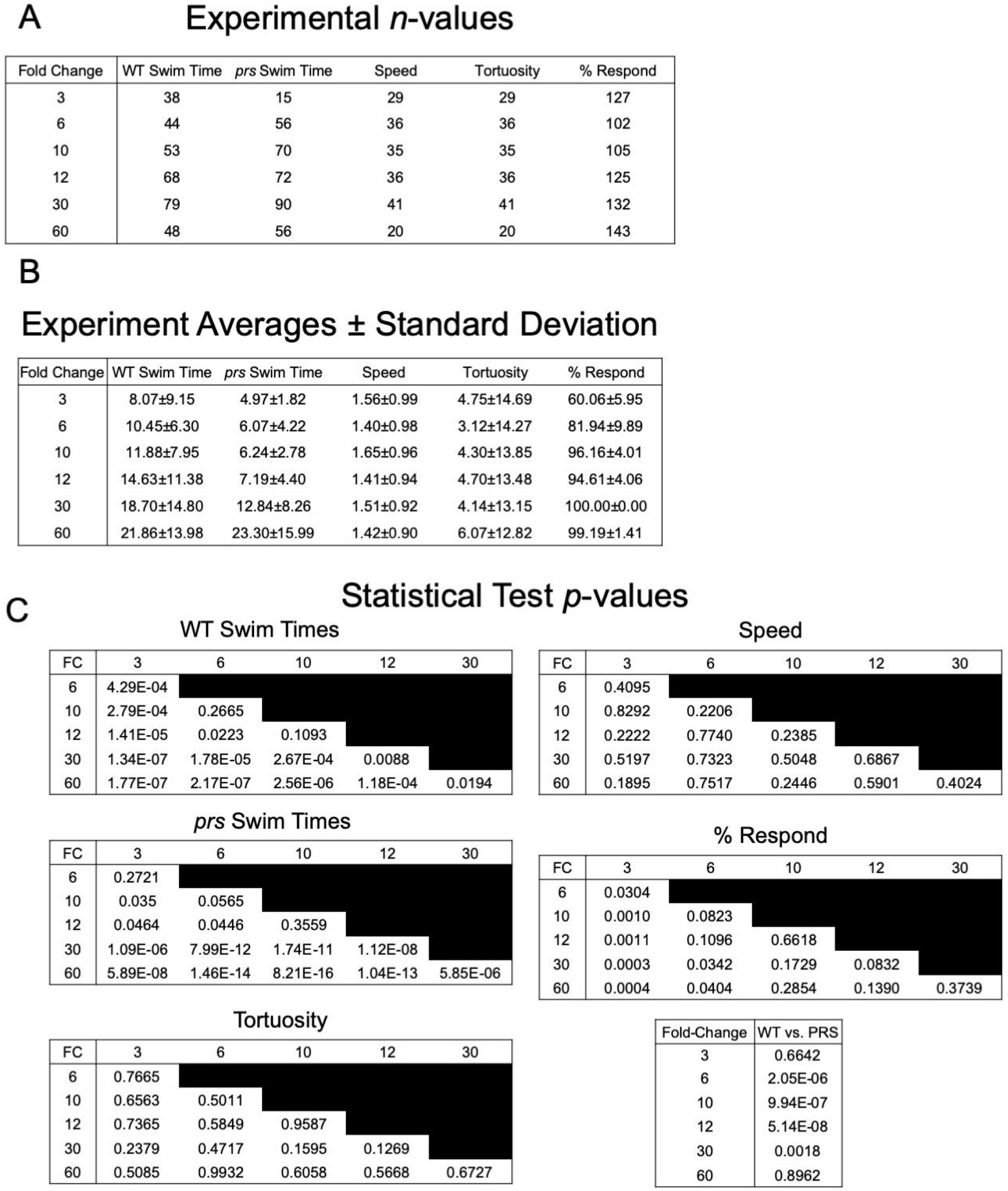
Fold Change Detection Behavior. **A**. Number of larvae (n) assessed in the fold change series (3 - 60, first column) for WT swim time (panel 2A), speed (panel 2B), tortuosity (panel 2C), percent responding (panel 2D), and prs swim time (panel 2F). **B**. The experimental averages and standard deviations for the assays described in panel A. **C.** Pairwise tests for significance at each fold-change for swim time (WT and *prs*), speed, percent response and tortuosity (p-values shown; all comparisons by Wilcoxon, except percent response, which was done by T-test). Also shown (bottom right) is a comparison of swim times in WT and *prs* larvae at each fold-change step. p-values for each comparison are listed. WT = wild type; *prs* = *pristine* mutant; FC = fold change.

**Figure 3 - Source Data 1.**
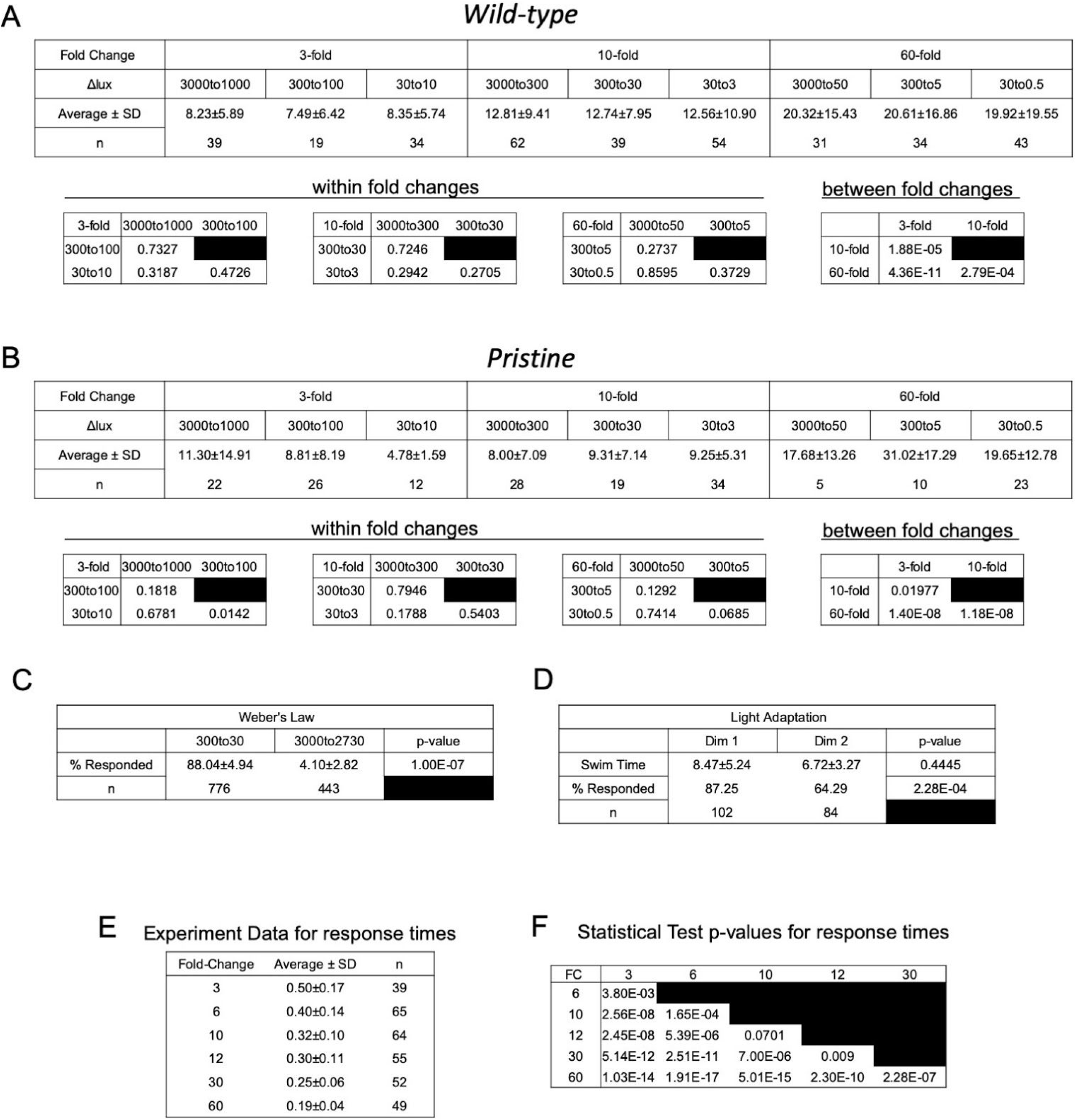
Validation of fold change detection behavior. **A.** Average swim times, standard deviations, and number of larvae analyzed for data presented in Figure 3A. **B**. Average swim times, standard deviations, and number of larvae analyzed for data presented in Figure 3B. **C**. Data for results plotted in Figure 3C. Sample size (n), percent of larvae responding, and test of significance (p-value; T-test) are shown. **D**. Data for results plotted in Figures 3E and F. Included in the table are the number of larvae analyzed at each dim, the average swim times (and standard deviations). Statistical analyses indicate that fewer larvae responded to Dim 2, but the average swim times were not different. **E.** Average response times, standard deviations, and number of larvae analyzed for data presented in Figure 3G. **F.** Pairwise statistical analysis of response times for results shown in Figure 3G. (p-values shown; all comparisons by Wilcoxon).

**Figure 5 - Source Data 1.**
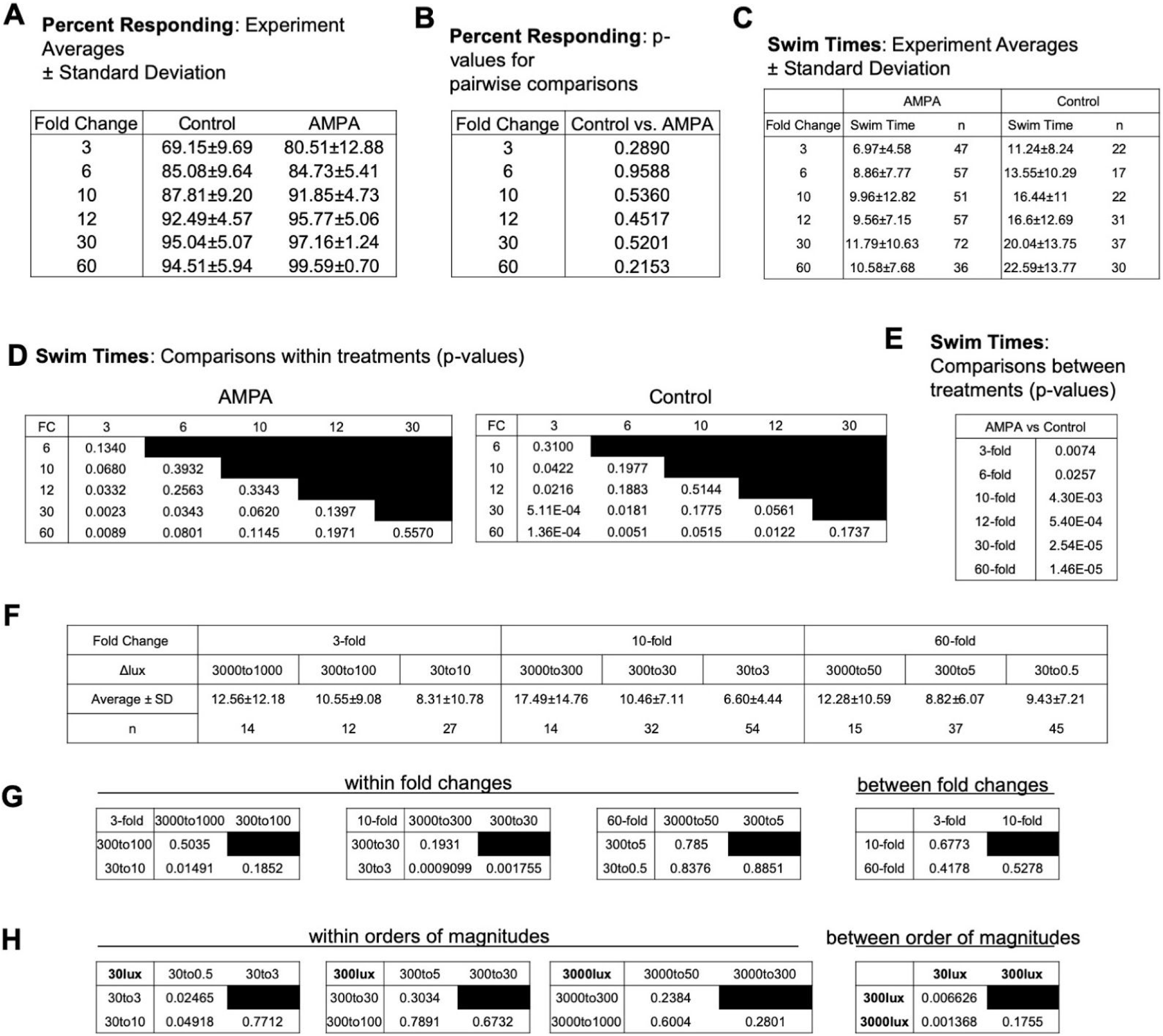
AMPA disrupts fold change detection. **A.** Averages of percent of control and AMPA treated larvae responding to the indicated fold dimming for results plotted in Figure 5B. **B**. p-values (T-test) for pairwise comparisons at each fold change for results plotted in Figure 5B. **C.** Sample sizes (n), average swim times and standard deviations for data plotted in Figure 5C. **D**. Statistical analyses for data plotted in Figure 5C. All pairwise comparisons were made within treatments (AMPA or control) for each fold change (FC). p-valves are shown (Wilcoxon). **E.** Comparison of swim times at each fold change between treatments for results plotted in Figure 5C. p-valves are shown (Wilcoxon). **F.** Sample sizes (n), averages and standard deviations for results plotted in Figure 5D and E. **G**. Statistical analyses of swim times of AMPA-treated larvae for data plotted according to fold change (Figure 5D). All tests within a fold change group, and between fold changes, was performed by Wilcoxon (p-values shown). **H.** Statistical analyses of swim times of AMPA-treated larvae for data plotted according to initial light intensity (Figure 5E). The initial light intensity (before dim) and are indicated as 30lux, 300lux and 300lux in the figure. All tests within an initial intensity group, and between initial intensities, was performed by Wilcoxon (p-values shown).

